# Asteroid: a new minimum balanced evolution supertree algorithm robust to missing data

**DOI:** 10.1101/2022.07.22.501101

**Authors:** Benoit Morel, Tom A. Williams, Alexandros Stamatakis

**Affiliations:** Computational Molecular Evolution group, Heidelberg Institute for Theoretical Studies, Heidelberg, Germany; Institute for Theoretical Informatics, Karlsruhe Institute of Technology, Karlsruhe, Germany; School of Biological Sciences, University of Bristol, Bristol, UK

## Abstract

**Motivation:** Missing data and incomplete lineage sorting are two major obstacles to accurate species tree inference. Gene tree methods such as ASTRAL and ASTRID have been developed to account for incomplete lineage sorting. However, they can be severely affected by high levels of missing data.

**Results:** We present Asteroid, a novel supertree method that infers an unrooted species tree from a set of unrooted gene trees. We show on both empirical and simulated datasets that Asteroid is more robust to missing data than ASTRAL and ASTRID, while being several orders of magnitude faster than ASTRAL for datasets that contain thousands of genes. Asteroid offers advanced features such as parallelization, support value computation, and support for multi-copy and multifurcating gene trees.

**Availability:** Asteroid is freely available at https://github.com/BenoitMorel/Asteroid

**Contact:** online.

## 1 Introduction

Due to recent advances in sequencing technology, biologists now have access to an exponentially increasing quantity of phylogenomic data that can help to unravel evolutionary relationships between species. However, despite this abundance of informative data, inferring species trees remains a challenging task, especially when speciation events occured within very short time intervals.

An important obstacle to accurate phylogenetic tree inference is the discordance between species and gene histories. Biological processes such as incomplete lineage sorting (ILS), gene duplication, gene loss, or horizontal gene transfer, yield species trees that are topologically distinct from the respective gene trees. In particular, it has been shown that concatenation approaches (Minh *et al*., 2020; Kozlov *et al*., 2019; Lartillot *et al*., 2009; Ronquist *et al*., 2012; Aberer *et al*., 2014) (also called supermatrix methods) are inconsistent under the multi-species coalescent (MSC) model (Kubatko and Degnan, 2007), which is widely used to model ILS (Rannala and Yang, 2003).

To this end, supertree methods have been developed to better accommodate the incongruence between the gene trees and species history. They initially infer one gene tree per gene sequence alignment, and subsequently estimate the species tree from the inferred set of input gene trees. Some of these methods are particularly appealing because they can process large datasets and are consistent under the MSC model. For instance, ASTRAL-III (Zhang *et al*., 2018) defines the quartet score of a species tree as the number of quartets induced by the input gene trees that are compatible with the species tree. ASTRAL-III implements a search heuristic that first extracts a list of all bipartitions from the input gene trees, and subsequently deploys a dynamic programming algorithm to traverse all species trees whose bipartitions are contained in this list. ASTRAL-III computes the quartet score for each of those candidate species trees, and returns the tree with the best quartet score. FastRFS (Vachaspati and Warnow, 2016) uses the same exploration strategy to search for the species tree. In contrast to ASTRAL-III, it attempts to minimize the Robinson-Foulds (RF) distance (Robinson and Foulds, 1981) between the species tree and the gene trees. The search strategy used by ASTRAL-III and FastRFS allows them to analyze datasets comprising a large number of species. However, their runtime complexity is more than quadratic with respect to the number of genes (*O*((*NK*)^2.726^), for ASTRAL-III, and *O*((*NK*)^3^) for FastRFS, where *N* is the number of species, and *K* the number of genes), and they therefore become computationally demanding with increasing number of genes. ASTRID (Vachaspati and Warnow, 2015) is a faster alternative that runs in time that is linear in the number of genes (*O*(*KN* ^3^)). It defines the internode distance between two leaves in a tree as the number of internal nodes on the path connecting those two leaves. ASTRID computes a distance matrix whose elements represent the distance between each pair of species, as estimated from the gene tree leaf internode distances. Then, it estimates a species tree from this matrix under the minimum balanced evolution principle (Lefort *et al*., 2015) (see our Method section for a more detailed description of ASTRID). ASTRAL-III, FastRFS, ASTRID, as well as other supertree methods, are consistent under the MSC model, assuming that the gene trees have been correctly estimated *and* that each gene tree comprises all species.

However, the consistency of the above methods is not well established for datasets with missing data, which represent the standard use case in biological practice. In the following, as *missing data*, we consider gene trees that do not comprise all species under study. Missing data can have various causes, such as gene loss events, errors in the gene orthology clustering process, or genes simply not being sequenced. In addition, some pre-processing pipelines filter out gene sequences, for instance by identifying abnormally long gene tree branches (Mai and Mirarab, 2018), by removing sequences with too many gaps (Wickett *et al*., 2014), or by identifying and removing rogue gene sequences (Aberer *et al*., 2013; Chen *et al*., 2021). Missing data can be considered as the result of a stochastic process that, starting from a comprehensive gene tree comprising all species, randomly deletes gene leaves. We refer to the models describing this stochastic process as *models of taxon deletion*. It has been shown that under some very specific models of taxon deletion, ASTRAL-I and ASTRAL-II are still consistent under the MSC model (Nute and Chou, 2017), but that ASTRID is inconsistent (Rhodes *et al*., 2020). However, the consistency of ASTRAL as well as other methods such as FastRFS has not yet been established for more general models of taxon deletion.

Here, we show that widely used methods such as ASTRAL-III, FastRFS, and ASTRID are not robust to high proportions of missing data. To address this issue, we introduce a new algorithm and open-source implementation, Asteroid, that constitutes a modification of the ASTRID algorithm and can better account for missing data. To achieve this in Asteroid, we introduce a new optimization criterion, the *global induced length* of a tree. We show that this optimization criterion is consistent under any model of taxon deletion. Our experiments on simulated datasets show that Asteroid is as accurate as the competing supertree methods in the absence of missing data, and considerably more accurate for large proportions of missing data. Further, we provide several empirical examples where Asteroid infers biologically plausible trees in contrast to the competing methods that fail to recover widely accepted and well-established phylogenetic relationships. In addition, on these empirical datasets, Asteroid is two orders of magnitude faster than ASTRAL-III and three orders of magnitude faster than FastRFS. It is, in the worst case, only one order of magnitude slower than ASTRID. Asteroid provides several useful features, such as bootstrap support value computation, conducting tree searches from multiple starting trees, support for multifurcating as well as multi-copy gene trees, and a distributed memory parallelization. Finally, we evaluated the quality of our implementation with respect to coding standards adherence using the SoftWipe (Zapletal *et al*., 2021) tool. The SoftWipe score of Asteroid is 8.6*/*10, which places Asteroid second in the list of 52 scientific tools written in C/C++ that the SoftWipe benchmark comprises.

## 2 Method

We first introduce some necessary notations and definitions. Then, we review the definition of the *global length* of a species tree (Lefort *et al*., 2015; Vachaspati and Warnow, 2015). We then explain our extension to define the *global induced length* of a tree. Subsequently, we outline our new method, Asteroid and how it efficiently searches for the species tree that minimizes our global induced length.

### 2.1 Preliminaries

Let *T* be an unrooted binary tree with |*T* | taxa. The *internode distance* between two taxa *i* and *j* of *T* is the number of internal/inner nodes on the path connecting *i* with *j* in *T*. Let *L*(*T*) denote the taxon set of *T*. Further, let *X* ⊂ *L*(*T*). The tree *T*_|*X*_ *induced* by *T* and *X* is the tree obtained from *T* by removing all taxa that are not contained in *X* and by subsequently contracting all nodes of degree one until the tree becomes strictly binary. A *species tree S* is an unrooted binary tree. Given a species tree *S*, a *gene tree* is an unrooted tree such that *L*(*G*) ⊂ *L*(*S*). Note that a gene tree *can* be multifurcating. *L*(*G*) represents the set of species taxa covered by the gene tree *G* and is called the *coverage pattern* of *G*.

### 2.2 Global length of a species tree in ASTRID

Let *S* be a species tree with *N* = |*S*| taxa. Let *M*_*S*_ be the matrix whose elements *M*_*S*_(*i, j*) represent the pairwise internode distances between the taxa of *S*. Let G = (*G*_1_, *G*_2_, …, *G*_*K*_) be a set of *K* gene trees. Let *F* (*i, j*) be the set of indices of the gene trees that comprise taxa *i and j*. Let *D*_*k*_(*i, j*) be the internode distance between the gene taxa *i* and *j* in *G*_*k*_ if *k* ∈ *F* (*i, j*), and 0 otherwise. Let *D*_G_ be the average internode distance matrix, such that for all taxon indices *i* and *j*, 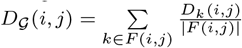.

The *global length* of *S* and G is defined as:

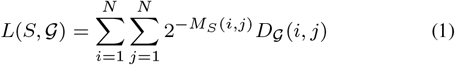

ASTRID (Vachaspati and Warnow, 2015) aims to find the species tree *S* that minimizes this global length. Note that the absence of a set of taxa in a gene tree might result in underestimating the internode distance between the remaining species that *are* present in this gene tree. As a result, including genes with missing data introduces a bias that will, in some cases, mislead the species tree inference process. This even holds, in the absence of ILS and in case the gene trees are correctly inferred (see Fig. 1).

**Fig. 1:**
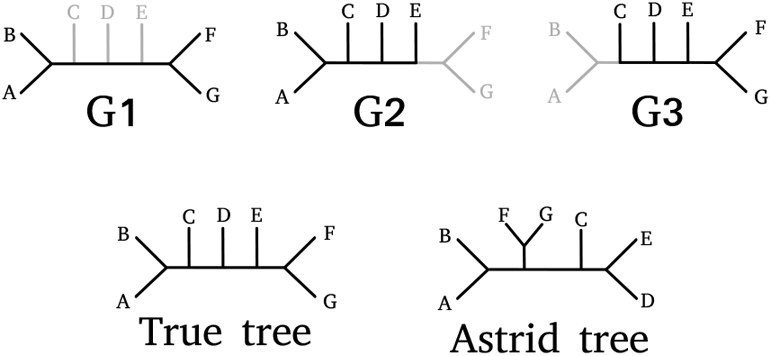
Example of the missing data bias causing ASTRID to infer an incorrect species tree. *G*_1_, *G*_2_, and *G*_3_ are three incomplete gene trees that perfectly agree with the true species tree (“True tree”) shown. Grey lines correspond to non-sampled subtrees. The gene tree *G*_1_ is the only tree that covers both species *A* and *F*. The internode distance between *A* and *F* is 5 in the true species tree, but only 2 in *G*_1_. Similarly, the internode distances between *A* and *G*, between *B* and *F*, and between *B* and *G* are all being underestimated because of missing genes. This introduces a bias in the resulting distance matrix, placing the clades (*A, B*) and (*F, G*) closer to each other than they should be.

### 2.3 Global induced length of a species tree

We introduce the *global induced length* of a tree to correct for this missing data bias. Instead of merging all gene tree internode matrices *D*_*k*_ into one single matrix *D*_G_, we separately compute the global length for every gene tree and matrix. We then apply an appropriate correction based on the respective taxon coverage patterns.

For each 1 ≤ *k* ≤ *k*,let 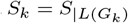 be the species tree induced by *S* and the set of species covered by the gene tree *G*_*k*_. Let *M*_*k*_ be the matrix whose elements contain the pairwise internode distances between the taxa of *S*_*k*_. The *global induced length* is defined as:

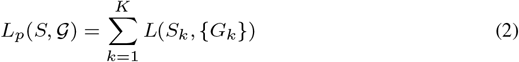

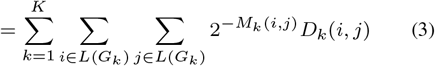

Separately weighting each gene tree internode distance matrix *D*_*k*_ with the corresponding induced species tree distance matrix *M*_*k*_ (instead of *M*) corrects for the gene internode estimation bias and yields the global induced length more robust to missing data. In the Supplementary Material, we prove that the global induced length is consistent under the multi-species coalescent model and under any random model of taxon deletion that is independent across genes, under the assumption that the gene trees have been correctly estimated. In other words, when the number of input gene trees increases asymptotically, the global induced length is minimized by the true species tree.

### 2.4 Asteroid

Asteroid is a search heuristic that aims to find the species tree with the lowest global induced length. In this section, we briefly outline our approach. A more detailed description of our algorithm can be found in the Supplementary Material.

Asteroid starts from a given species tree topology and applies a tree search strategy based on subtree prune and regraft (SPR) moves to further improve the score. The starting species tree can be either a random tree or a tree estimated with our own re-implementation of the ASTRID algorithm. Then, we iteratively optimize the species tree topology by adapting the FastME tree search algorithm (Lefort *et al*., 2015) to the global induced length score. Note that, at each step, the original FastME algorithm evaluates all possible SPR moves, but only applies the best one. To accelerate convergence, Asteroid simultaneously applies all non-conflicting SPR moves that improve the global induced length.

### 2.5 Multi-copy and multifurcating gene trees

Asteroid supports both, multi-copy, and multifurcating input gene trees.

When a gene tree covers the same species strictly more than once, we compute the corresponding gene tree distance matrix using the MiniNJ approach (Morel *et al*., 2022): the distance between two species is the minimum internode distance among all gene leaves that are mapped to those two species.

Accounting for polytomies in the gene trees does not require any change in the Asteroid algorithm, because the internode distance is also well defined for multifurcating gene trees.

### 2.6 Support value estimation

To assess confidence in the inferred species tree, Asteroid computes branch support values via multi-locus bootstrapping (Seo, 2008). Let *K* be the number of gene trees used to infer the best scoring tree, let *m* be the user-given number of bootstrap species trees to compute, and let 𝒢_*b*_ be a set of user-provided bootstrap gene trees. Note that, 𝒢_*b*_ can also be set to G, the set of gene trees used to infer the best-scoring tree. At every iteration, Asteroid builds a bootstrap replicate by sampling *K* gene trees from 𝒢_*b*_ with replacement, and then computes a bootstrap species tree from this replicate. It repeats the procedure *m* times to infer *m* bootstrap species trees. Then, it applies the Felsenstein bootstrap algorithm (Felsenstein, 1985) to compute the branch support values on the best-scoring species tree.

## 3 Experiments

### 3.1 Tested methods

We compared Asteroid with three competing algorithms: ASTRAL-III, FastRFS, and ASTRID. We ran all the experiments on the same machine with 40 physical cores and 754GB of main memory. Asteroid and ASTRAL-III both offer a parallel implementation, but for a fair comparison, all reported runtimes were obtained by running the tools *without* parallelization in sequential mode. However, in experiments that do not focus on runtime performance, we occasionally executed the parallel versions of Asteroid and ASTRAL-III (ASTRAL-MP (Yin *et al*., 2019)) for the sake of convenience. Furthermore, the reported runtimes for Asteroid do not include bootstrap tree computations. We ran the FastME version of ASTRID with both NNI and SPR moves (-s option). We did not run the BioNJ version of ASTRID, which attempts to account for missing data, because the developers recommend not to use it (Vachaspati, 2021) and removed it from the tool. We ran Asteroid with the option -r 0 (which infers the starting tree with the ASTRID algorithm).

### 3.2 Dataset simulation

We generated simulated datasets with SimPhy (Mallo *et al*., 2015) to assess the influence of various model parameters on the reconstruction accuracy of the tested methods.

The parameters we studied are: the level of missing data, the level of discordance due to ILS, the number of gene trees, the number of species, the average sequence length, and the *gene tree branch length scaler*, which is a multiplication factor that we use to rescale the gene tree branch lengths. The level of discordance due to ILS is defined as the average RF distance between the true gene trees and the true species tree *before* deleting genes.

For each gene, we sampled the sequence length from a log-normal distribution (with, by default, 100 sites per sequence on average).

To simulate missing data, we randomly sampled gene sequences from the simulated datasets. In a first step, we assigned to each species *a* a deletion probability *d*_*s*_(*a*) and to each gene *k* a deletion probability *d*_*f*_ (*k*). In a second step, we deleted every gene sequence belonging to gene *k* and species *a* from the initial dataset with probability *d*_*s*_(*a*)*d*_*f*_ (*k*). When a species was not covered by any gene, we removed it from the reference species tree. When a gene contained less than 4 gene sequences, we excluded it from the dataset. We sampled the probabilities *d*_*s*_(*a*) and *d*_*f*_ (*k*) from a Beta distribution, because this distribution yields values between 0 and 1, allows for generating various and highly distinct distribution shapes, and matches our observations on empirical missing data patterns, as we show in the supplementary material. In order to better control the average proportion of missing data, we used a re-parametrization *B*^∗^(*μ, θ*) of the standard Beta distribution *B*(*α, β*), where 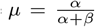 is the mean and *ν* = *α* + *β* the sample size of the distribution.

We inferred the gene trees with ParGenes (Morel *et al*., 2018), performing one RAxML-NG maximum likelihood search from a single random starting tree per gene under the general time reversible model of nucleotide substitution with four discrete gamma rates (GTR+G4) (Tavaré *et al*., 1986; Yang, 1993). Then, we inferred the species trees with all tested tools from this set of inferred gene trees.

### 3.3 Effect of missing data distribution

In this first experiment, we studied how missing data affects the tested methods in absence of ILS. We selected a default set of simulation parameters and applied different random distributions of taxon deletion. The default parameters are: 50 species, 200 genes, 100 sites per sequence (on average), and no ILS. We provide a more detailed description of the simulation parameters in the Supplementary Material.

We first studied the effect of missing data under heterogeneous deletion probabilities across *species*. We set *d*_*f*_ (*k*) := 0 (no per-partition deletion) and drew the per-species deletion probabilities *d*_*s*_(*a*) from *B*^∗^(*μ*_*s*_, 5) for different values of *μ*_*s*_.

Then, we studied the effect of missing data under heterogeneous deletion probabilities across *genes*. We set *d*_*s*_(*a*) := 0 and drew the per-gene deletion probability *d*_*f*_ (*k*) from *B*^∗^(*μ*_*f*_, 5) for different values of *μ*_*f*_.

Finally, we studied the effect of missing data under heterogeneous deletion probabilities across both, genes, and species. We drew both *d*_*f*_ (*k*) and *d*_*s*_(*a*) from *B*^∗^(*μ*, 5) for different values of *μ*, the mean deletion probability. Since we apply two deletion steps (per-species and per-gene), the resulting mean proportion of missing data is 1 − (1 − *μ*)^2^.

### 3.4 Effect of remaining simulation parameters

In this second experiment, we studied the effect of the remaining simulation parameters in the presence of ILS for high proportions of missing data. We defined a default set of simulation parameters, and varied each parameter individually. The default parameters are: 50 species, 1000 genes, 100 sites per sequence (on average), and a population size of 5, 0000, 000 (average level of discordance: 0.22). The parameters that we varied were the number of species, the average sequence length, the average proportion of missing data, the population size, the number of genes, and the gene tree branch length scaling factor. We provide a detailed list of the simulation parameters in the Supplementary Material. By default, *μ*_*f*_ = *μ*_*s*_ := 0.6, which means that we first removed 60% of the gene sequences under heterogeneous deletion probabilities across species, and then 60% of the remaining gene sequences under heterogeneous deletion probabilities across partitions.

### 3.5 Empirical datasets

We studied the performance of the tested methods on three different phylogenomic datasets: a plant dataset with 81 species, a vertebrate dataset with 179 species, and a dataset with 92 Eukaryote and Archaea species. We generated two versions of each of these three datasets: the *single* datasets only contain the initial single-copy gene trees. We generated the *disco* datasets by inferring or extracting multi-copy gene trees, and subsequently decomposing them into single-copy gene trees using DISCO (Willson *et al*., 2021). We only kept the gene trees covering at least four different species. Table 1 summarizes the dimensions of the resulting six datasets. Finally, in order to study the effect of an increasing proportion of missing genes, we filtered out the most gappy gene sequences from the vertebrates single-copy gene alignments using different threshold values.

**Table 1.**
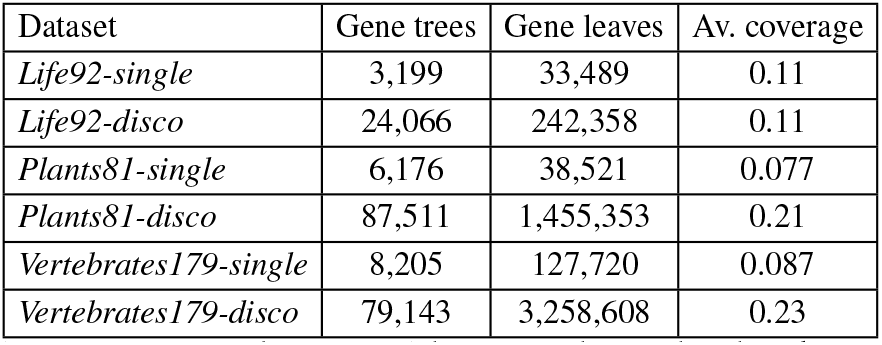
Description of the empirical datasets used in our benchmark. Dataset names are suffixed by the number of species in the respective dataset. Gene trees is the number of gene trees. Gene leaves is the total number of gene tree leaves. Av. coverage is the average number of species covered by each each gene tree.

| Dataset | Gene trees | Gene leaves | Av. coverage |
| --- | --- | --- | --- |
| *Life92-single* | 3,199 | 33,489 | 0.11 |
| *Life92-disco* | 24,066 | 242,358 | 0.11 |
| *Plants81-single* | 6,176 | 38,521 | 0.077 |
| *Plants81-disco* | 87,511 | 1,455,353 | 0.21 |
| *Vertebrates179-single* | 8,205 | 127,720 | 0.087 |
| *Vertebrates179-disco* | 79,143 | 3,258,608 | 0.23 |

#### 3.5.1 Eukaryote and Archaea dataset

The dataset *Life92-single* consists of 3199 single-copy gene trees covering 92 species from the Eukaryote and Archaea domains. The corresponding gene trees were inferred in a previous study (Williams *et al*., 2020). We extracted the *Life92-disco* dataset from 41,222 multi-copy gene trees that we had also inferred in another study (Morel *et al*., 2022). We applied DISCO to those gene trees to obtain 24,066 single-copy gene trees. The number of single-copy gene trees is lower than the number of input multi-copy gene trees because we filtered out the gene trees that comprised less than four distinct species.

#### 3.5.2 Plant dataset

The dataset *Plants81-single* consists of 6,176 single-copy gene trees covering 81 plant species. We extracted the corresponding gene multiple sequence alignments from release 59 of the Ensembl Plants database (Bolser *et al*., 2016). Then, we inferred the gene trees with ParGenes, performing one RAxML-NG maximum likelihood search from a single random starting tree per gene under the LG+G model (Le and Gascuel, 2008). We also extracted 51,461 multi-copy gene sequence alignments from the same database, and inferred the corresponding gene trees with ParGenes, again performing one RAxML-NG maximum likelihood search from a single random starting tree per gene under the LG+G model. We obtained the *Plants81-disco* dataset by decomposing these multi-copy gene trees into 87,511 single-copy gene trees with DISCO.

#### 3.5.3 Vertebrate dataset

The dataset *Vertebrates179-single* consists of 8,205 gene trees covering 179 vertebrate species. We extracted the single-copy multiple sequence alignments from release 105 of the Ensembl Compara database (Zerbino *et al*., 2017). We also generated several filtered datasets, by removing gene sequences with a proportion of gaps exceeding a threshold value *τ*, for different values of *τ*. The filtered datasets are described in Table 2. We also extracted all 33, 809 multi-copy sequence alignments from the Ensembl database, inferred the corresponding gene trees, and applied DISCO to obtain the *Vertebrates179-disco* dataset with 79,143 single-copy gene trees. We inferred all gene trees (from the single-copy, multi-copy, and filtered alignments) with ParGenes, performing one RAxML-NG maximum likelihood search from a single random starting tree per locus under the GTR+G model.

**Table 2.**
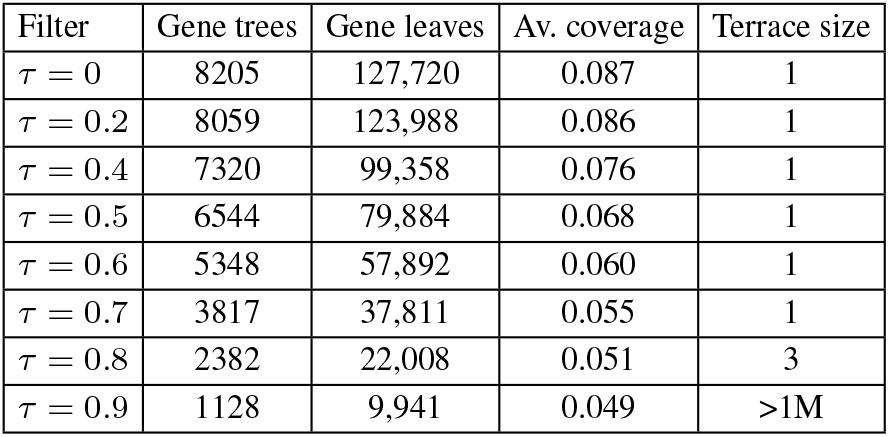
Dimensions of the datasets obtained by filtering gene sequences of the Vertebrates179-single that exhibit a proportion of non-gap characters below *τ* (*τ* = 0 corresponds to the unfiltered dataset). Gene trees is the number of gene trees. Gene leaves is the total number of gene tree leaves (summed over all the gene trees). Average coverage is the average number of species covered by each each gene tree. Terrace size is the number of species trees that have the same global induced length as the Asteroid tree (see the Supplementary Material for more details about terrace computation).

| Filter | Gene trees | Gene leaves | Av. coverage | Terrace size |
| --- | --- | --- | --- | --- |
| $\tau = 0$ | 8205 | 127,720 | 0.087 | 1 |
| $\tau = 0.2$ | 8059 | 123,988 | 0.086 | 1 |
| $\tau = 0.4$ | 7320 | 99,358 | 0.076 | 1 |
| $\tau = 0.5$ | 6544 | 79,884 | 0.068 | 1 |
| $\tau = 0.6$ | 5348 | 57,892 | 0.060 | 1 |
| $\tau = 0.7$ | 3817 | 37,811 | 0.055 | 1 |
| $\tau = 0.8$ | 2382 | 22,008 | 0.051 | 3 |
| $\tau = 0.9$ | 1128 | 9,941 | 0.049 | >1M |

We generated a multifurcating reference species tree by starting from the NCBI tree of the 179 species, and contracting the splits that were contradicted by at least one of the tested tools *and* by several independent studies. We describe the five splits that we contracted in the supplementary material.

## 4 Results

### 4.1 Effect of missing data in absence of ILS

We summarize the results of the simulated data experiment without ILS in Fig. 2. We first remark that, for the same mean proportion of missing data, all methods are substantially more sensitive to heterogeneous taxon deletion probabilities across species than across genes. For instance, for simulations with 80% of missing data, the average RF distance between the Asteroid species trees and the true species tree is 0.23 when all species have the same average coverage, and 0.55 when species have a different average coverage. Among-taxa variation in the proportion of missing data is common in empirical datasets because of species- or clade-specific differences in patterns of genome evolution, such as elevated rates of gene loss and genome reduction in host-associated microbes (McCutcheon and Moran, 2012). This result indicates that such patterns are likely to be particularly challenging for accurate species tree inference. For all simulation parameters in this first experiment, we observe the same trend: all tested methods perform analogously for low proportions of missing data (less than 50%). For high proportions of missing data, Asteroid performs better than both ASTRAL-III and FastRFS, and ASTRID performs considerably worse than all other methods. For instance, for the highest proportions of missing data with heterogeneous rates across both species and genes, Asteroid finds the most accurate trees (*RF* = 0.51), followed by FastRFS (*RF* = 0.59), ASTRAL-III (*RF* = 0.61), and ASTRID (*RF* = 0.91).

**Fig. 2:**
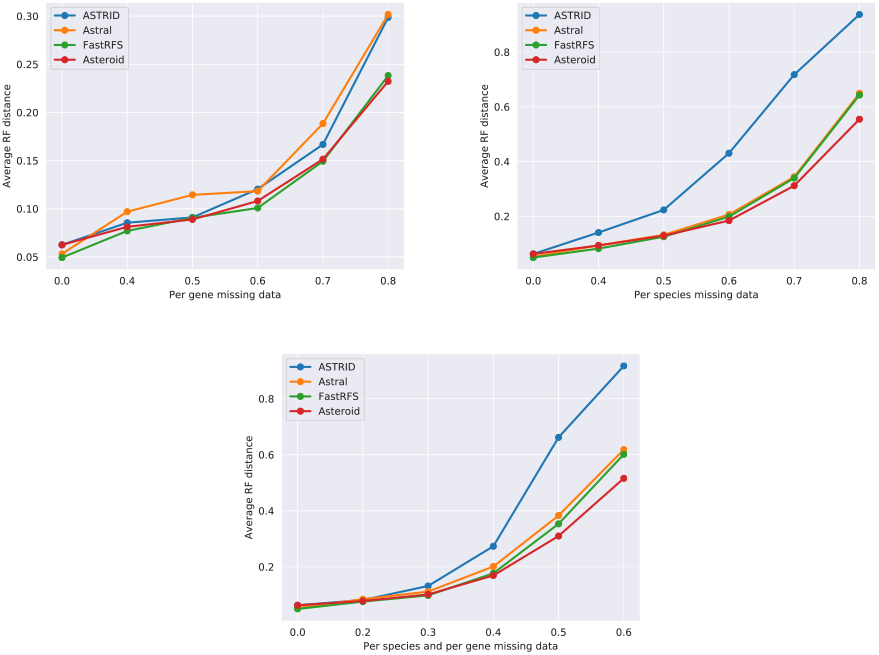
Average unrooted RF distance between inferred and true species trees in absence of ILS for varying proportions of missing data

**Fig. 3:**
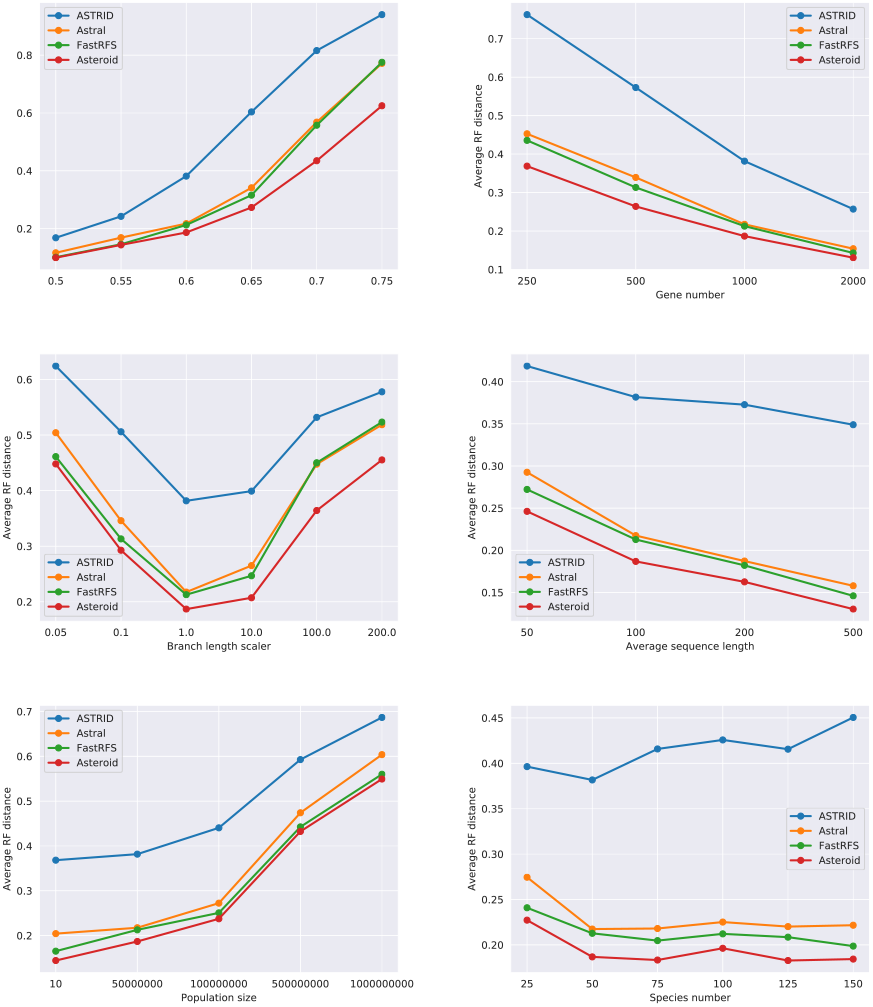
Average unrooted RF distance between inferred and true species trees for high levels of missing data.

### 4.2 Effect of various parameters under high proportions of missing data

In the second experiment, we observe the same trend: Asteroid is the most accurate method under most simulation parameters. FastRFS and ASTRAL-III perform similarly, with a small advantage for FastRFS. ASTRID is considerably less accurate. The difference between the methods increases for increasing proportions of missing data, until the RF distance becomes saturated. All tools achieve better performance when the number of genes increases and when the gene tree reconstruction error decreases. The gene tree discordance due to ILS negatively affects reconstruction accuracy for all methods, but affects all of them similarly. Finally, the number of species does not appear to affect the average RF distance to the true tree.

### 4.3 Plants81 biological dataset

On the Plants81-single dataset, Asteroid recovers a tree that is in good agreement with the reference taxonomy and a series of recent studies (Wickett *et al*., 2014; Zeng *et al*., 2014; Puttick *et al*., 2018; Initiative, 2019; Harris *et al*., 2020), with the exception of three specific branches.

First, it places Rosales within Malvids instead of within Fabids. Second, it places *Daucus carota* within Ericales instead of Campanulids. Finally, it places the Streptophyta clade between the two Chlorophyta species (*Chlamydomonas* and *Ostreococcus*), which traditionally form a monophyletic clade. All these conflicts are local (i.e., they only affect one branch) and the remaining splits are in agreement with the taxonomy. ASTRID, ASTRAL-III, and FastRFS all fail to infer a plausible tree and contradict the taxonomy tree on numerous and diverse splits that we do not list here. The RF distance to the NCBI taxonomy is 0.09 for Asteroid, 0.30 for ASTRAL-III, 0.37 for FastRFS, and 0.65 for ASTRID. We also attempted to run ASTRAL-III by extending its search space by the bipartitions of the species tree inferred with Asteroid, resulting in another tree with a higher quartet scores, and a RF distance to the taxonomy of 0.1.

On the Plants81-disco dataset, both Asteroid and ASTRID find highly similar trees, in agreement with recent literature (Wickett *et al*., 2014; Zeng *et al*., 2014; Puttick *et al*., 2018; Initiative, 2019; Harris *et al*., 2020). The Asteroid tree is presented in Fig. 4. The tree inferred by ASTRAL-III and FastME contradict the literature on one split among red alga, placing Chondrus crispus (Gigartinales) between Galdieria sulphuraria and Cyanidioschyzon merolae (Cyanidiales). On the disco dataset, all tested methods recover one relationship that, based on the literature (Wickett *et al*., 2014; Zeng *et al*., 2014; Puttick *et al*., 2018; Initiative, 2019; Harris *et al*., 2020), is unlikely to be correct: they recover the lycophyte *Selaginella* as sister to the bryophytes, rather than as sister to the euphyllophytes (angiosperms and ferns). This result might reflect the sparse sampling of bryophytes and lycophytes in the Plants81 dataset.

**Fig. 4:**
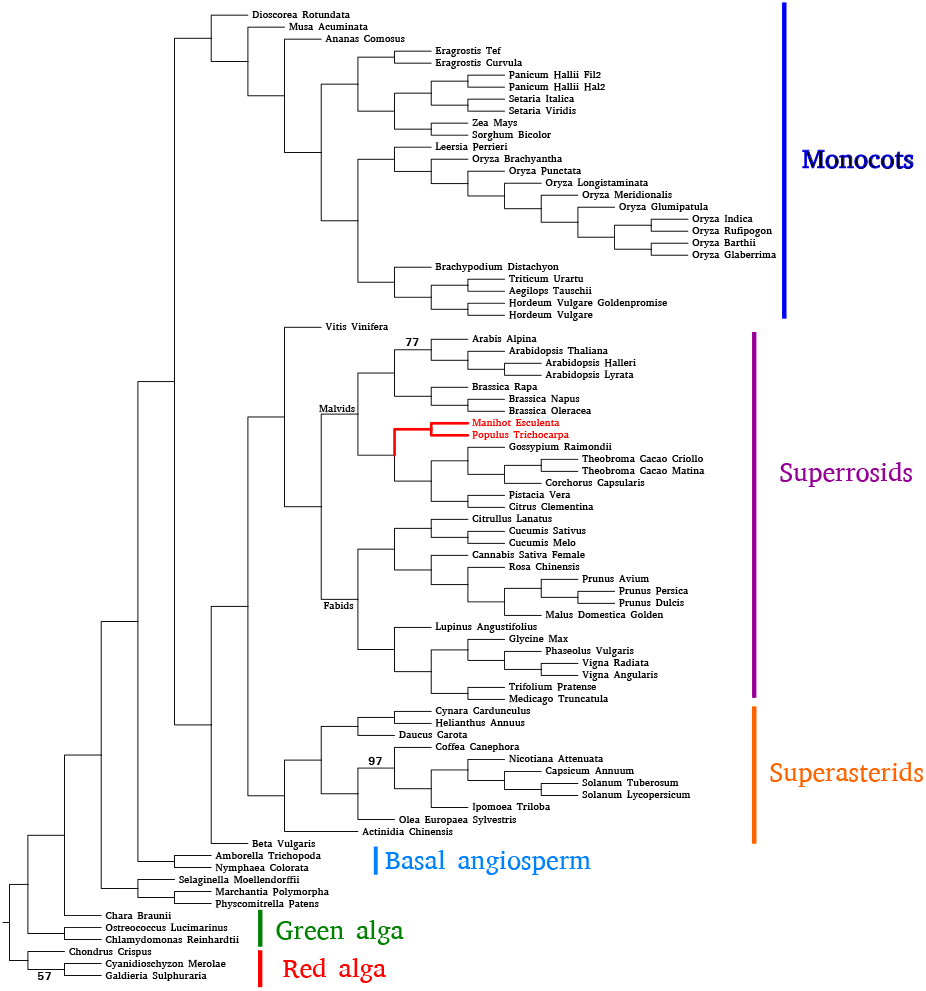
Species tree inferred with Asteroid from the Plants81-disco dataset. We placed the root manually. We only show support values below 100%. In red, we show the Malpighiales clade that is placed within Fabids in the NCBI taxonomy but within Malvids in all trees we inferred.

### 4.4 Life92 biological dataset

The Life92 dataset comprises representative genomes from the known major clades of eukaryotes and Archaea; some of the relationships among these groups are uncertain, while for others there is an emerging consensus in the literature. The Asteroid and FastRFS trees inferred from the Life92-single dataset are in good agreement with recent analyses using complex substitution models on protein concatenations (Williams *et al*., 2020; Liu *et al*., 2021). Based on the assumption that the tree is rooted elsewhere within the Archaea, Asteroid and FastRFS recover a sister-group relationship between Asgardarchaeota and eukaryotes (rather than a specific relationship between Heimdallarchaeota and eukaryotes (Williams *et al*., 2020)). The TACK (Thaumarchaeota, Aigarchaeota, Crenarchaeota and Korarchaeota) lineages are recovered as being monophyletic. Within eukaryotes, the metamonads are recovered as the group most distantly related to other eukaryotes, with the metamonad *Trimastix marina* branching between the other metamonads and the remaining eukaryotes (Williams *et al*., 2020).

The ASTRAL-III tree is similar to the Asteroid and FastRFS trees, but provides only weak support (0.16) for the monophyly of Asgardarchaeota; this does not imply that the method is less accurate, because the monophyly of Asgardarchaeota with respect to eukaryotes is currently uncertain (Williams *et al*., 2020; Liu *et al*., 2021). The ASTRAL-III and ASTRID trees fail to recover the monophyly of the SAR (Stramenopiles, Alveolates and Rhizarians) clade of eukaryotes, for which there is reasonable phylogenomic support (Burki *et al*., 2020). In addition, the ASTRID tree does not recover a specific relationship between Asgardarchaeota and eukaryotes, instead placing the Bathy- and Thaumarchaeota closest to the eukaryote stem in the unrooted tree.

Results with the Life92-disco dataset were broadly similar, with some exceptions. The Asteroid and FastRFS trees recover a clade of metamonads and discobans (monophyletic excavates), while ASTRAL-III and ASTRID support excavate paraphyly, with metamonads the earliest-diverging lineage within eukaryotes. ASTRAL-III fails to recover the monophyly of Asgardarchaeota, placing Odinarchaeota with the TACK+Euryarchaeota - a relationship that disagrees with published analyses (Zaremba-Niedzwiedzka *et al*., 2017; Williams *et al*., 2020; Liu *et al*., 2021), while ASTRID again fails to recover Asgardarchaeota as the closest relatives of eukaryotes in the unrooted tree.

### 4.5 Vertebrates179 biological dataset

On the Vertebrates179-single dataset, Asteroid, FastRFS, and ASTRAL-III perfectly agree with the reference tree. ASTRID contradicts the reference tree on four splits.

On the Vertebrates179-disco dataset, Asteroid, ASTRID, and ASTRAL-III all agree with the reference tree, except for one split: the elephant shark (*Chondrichtyian*), is placed within *Osteicthyians*, as sister to *Sarcopterygii*. This placement is likely to be incorrect because it contradicts the current consensus about elephant shark placement in the tree of life (Venkatesh *et al*., 2014). Interestingly, we found the same placement in another study (Morel *et al*., 2022) that compared several multi-copy gene supertree methods on the same dataset, suggesting a bias in the multi-copy gene trees. Due to excessive runtimes, we stopped FastRFS after one month of execution.

We then studied the effect of filtering out the gene sequences that exhibit a proportion of non-gap characters below *τ* for different values of *τ*. When *τ* increases, the number of deleted genes also increases, as we report in Table 2. As we show in Fig. 5, Asteroid is substantially more robust to this filtering strategy than the competing methods. It infers trees that are very close to the reference tree (normalized RF *<* 0.013) for values of *τ* ranging between 0.0 and 0.6, while all other methods start diverging from the reference tree at *τ* = 0.4. For instance, for *τ* = 0.6, the normalized RF distance to the true tree is 0.006 for Asteroid (one mismatching branch), 0.28 for ASTRAL-III and FastRFS, and 0.59 for ASTRID. The RF distance between the Asteroid and the true tree only starts to drastically increase for *τ* := 0.9 (*RF* = 0.36, against *RF* = 0.09 for *τ* = 0.8). This was expected, because the dataset with *τ* := 0.9 is indecisive with very high terrace sizes (more than 1, 000, 000 trees have the same optimal global induced length). Note that, when adding the Asteroid tree bipartitions to its search space, ASTRAL-III is able to recover the same tree as Asteroid for all filtered datasets. This again indicates, that the current ASTRAL-III search strategy might be sub-optimal under high proportions of missing data.

**Fig. 5:**
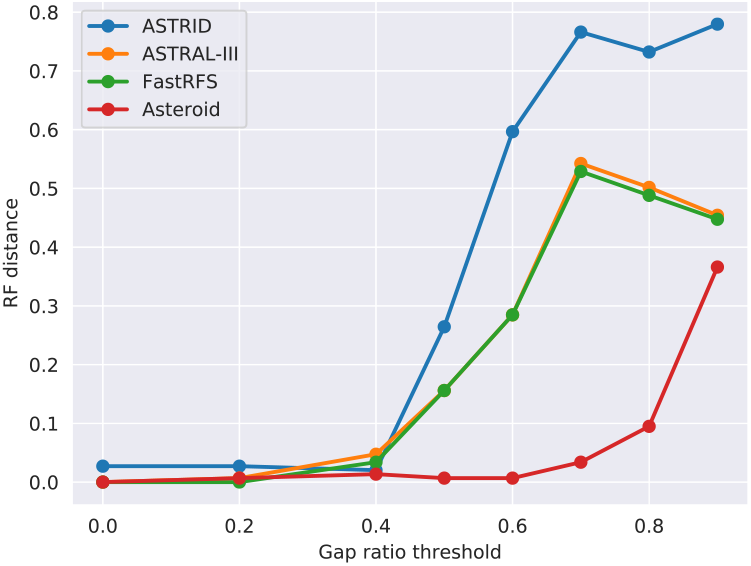
RF distance between the inferred and reference species tree for the dataset Vertebrates179 for different gap ratio thresholds.

### 4.6 Runtimes

We report the runtimes of the tested methods on the different empirical datasets in Table 3. ASTRID is the fastest method for most datasets. Asteroid is only four times slower than ASTRID in the worse case. On the disco datasets, Asteroid is more than two orders of magnitude faster than ASTRAL-III and three orders of magnitude faster than FastRFS.

**Table 3.**
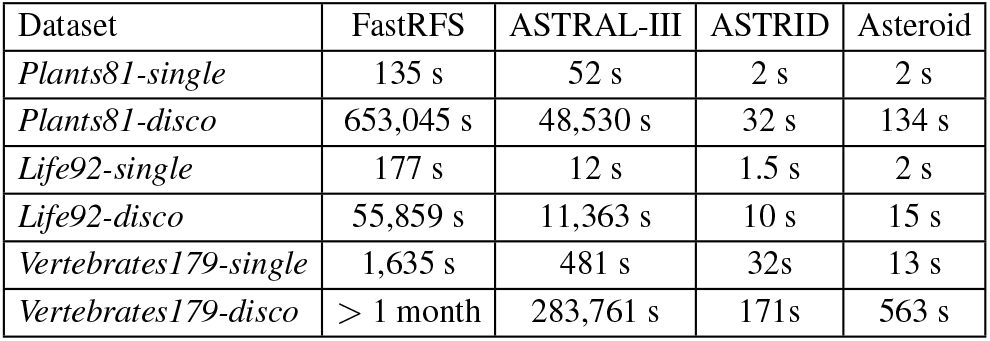
Sequential runtimes of the distinct methods on empirical datasets. Asteroid first estimates a starting tree with our implementation of the ASTRID algorithm and then applies the tree search algorithm presented in the Methods section). We terminated FastRFS on Vertebrates179-disco after one month.

| Dataset | FastRFS | ASTRAL-III | ASTRID | Asteroid |
| --- | --- | --- | --- | --- |
| *Plants81-single* | 135 s | 52 s | 2 s | 2 s |
| *Plants81-disco* | 653,045 s | 48,530 s | 32 s | 134 s |
| *Life92-single* | 177 s | 12 s | 1.5 s | 2 s |
| *Life92-disco* | 55,859 s | 11,363 s | 10 s | 15 s |
| *Vertebrates179-single* | 1,635 s | 481 s | 32s | 13 s |
| *Vertebrates179-disco* | > 1 month | 283,761 s | 171s | 563 s |

## 5 Discussion

Our experiments on both, simulated, and empirical data, suggest that Asteroid is more robust to missing data than the competing methods. In particular, ASTRID performs substantially worse than all other methods for large proportions of missing data.

The sub-optimal performance of ASTRID for high proportions of missing data (in comparison with other methods) is likely due to the missing data bias that we illustrate in Fig. 1: the values in the internode distance matrix computed by ASTRID underestimate the internode distance matrix of the true species tree. If the amplitude of this underestimation substantially differs from one pair of species to another, the distance matrix used by ASTRID will inevitably induce an incorrect species tree.

Surprisingly, our experiments also showed that Asteroid outperforms ASTRAL-III and FastRFS for large proportions of missing data. Unlike ASTRID, there is, to our knowledge, no reason to believe that the scores used by ASTRAL-III and FastRFS are biased in presence of missing data. Indeed, our proof that the global induced length is consistent under the MSC and any random of model of taxon deletion can also be adapted to the quartet score (ASTRAL-III) and to the RF distance between the gene trees and the species tree (FastRFS). However, an important difference between Asteroid and the two other methods is the heuristic used to explore candidate species trees. Both ASTRAL-III and FastRFS rely on the same search heuristic that consists in exploring all the species trees whose bipartitions are present in the input gene trees. Although ASTRAL-III deploys some strategies to accommodate incomplete gene trees when building the set of induced bipartitions (Zhang *et al*., 2018), we showed in the empirical experiments that ASTRAL-III might find a sub-optimal tree. Although there is no theoretical guarantee that it will find the best-scoring tree, the tree search heuristic implemented in Asteroid appears to be less affected by missing data, which might explain its good accuracy.

We note, however, that the differences between all tested methods decrease with increasing number of input gene trees. In particular, all methods inferred reasonable species trees for the three empirical datasets when using all the available data (by decomposing multi-copy gene trees into single-copy gene trees). We also observed the same trend under simulations: all methods find increasingly accurate species trees when the number of families increases, even under high proportions of missing data. Nonetheless, it is impossible to know how many gene trees are sufficient to alleviate the effects of missing data. In addition, it has been shown that, under specific conditions and for some models of taxon deletion, ASTRID is guaranteed to converge to an incorrect species tree when the number of gene trees increases asymptotically (Rhodes *et al*., 2020).

In terms of runtime, Asteroid has a clear advantage over ASTRAL-III and FastRFS for datasets with a large number of genes: it is up to two orders of magnitude faster than ASTRAL-III and up to three orders of magnitude faster than FastRFS. In particular, FastRFS failed to infer the Vertebrates179-disco species tree in less than one month, while Asteroid managed to complete within 10 minutes. ASTRID remains the fastest method and is at most one order of magnitude faster than Asteroid in our experiments, but at the cost of being inconsistent in the presence of ILS and missing data. It is also important to note that Asteroid can be run in parallel (unlike ASTRID and FastRFS), which might allow it to process even larger datasets in a reasonable amount of time.

## 6 Acknowledgments

This work was financially supported by the Klaus Tschira Foundation and by DFG grant STA 860/6-2. TAW is supported by a Royal Society University Fellowship. This work was funded by the Gordon and Betty Moore Foundation through grant GBMF9741 to TAW.

