## Supplement material for "Asteroid: a new minimum balanced evolution supertree algorithm robust to missing data"

picture(0,0)(-35,0)(1,0)30 (0,35)(0,-1)30 picture

bioRxiv picture(0,0)(35,0)(-1,0)30 (0,35)(0,-1)30 picture
doi/xxxxxx
Advance Access Publication Date: Day Month Year
Original Paper

Phylogenetics

### Supplement material for Asteroid: a new minimum balanced evolution supertree algorithm robust to missing data.

Benoit Morel,[1,2] Tom A. Williams,[3] and Alexandros Stamatakis,[1,2]

[1]Computational Molecular Evolution group, Heidelberg Institute for Theoretical Studies, Heidelberg, Germany
[2] Institute for Theoretical Informatics, Karlsruhe Institute of Technology, Karlsruhe, Germany
[3] School of Biological Sciences, University of Bristol, Bristol, UK

*To whom correspondence should be addressed.

Associate Editor: XXXXXXX

Received on XXXXX; revised on XXXXX; accepted on XXXXX

**Abstract**

Supplement material

#### 1 Consistency of the global induced length

In this section, we show that the global induced length is consistent under the multi-species coalescent model and under any random model of taxon deletion that is independent across genes, under the assumption that the gene trees have been correctly estimated. In other worlds, when the number of input gene trees increases asymptotically, the global induced length is minimized by the true species tree. Note however, that this true species tree might reside on a *terrace* in phylogenetic tree space (Sanderson *et al.*, 2015), that is, a set of species trees with the exactly the same analytical score. We wish to emphasize that the following proof does not induce that the Asteroid search heuristic will find the globally optimal tree.

Let $S$ be the ground-truth species tree. Let $\mathcal{C}_S$ be the set of all possible coverage patterns $C \subset L(S)$. A random model of taxon deletion $\mathcal{M}$ associates to each coverage pattern $C \in \mathcal{C}_S$ a probability $p(C)$, such that

$$\sum_{C \in \mathcal{C}_S} p(C) = 1$$

For a given such model of random taxon deletion $\mathcal{M}$, let $\mathcal{C}_\mathcal{M}$ be the set of coverage patterns $C$ with a non-zero probability $P(C)$. Let $\mathcal{G}_\mathcal{C}$ be the set of gene trees with the same coverage pattern $C$.

Lemma 1.1. *Let $C \in \mathcal{C}_\mathcal{M}$. When the size of $\mathcal{G}$ grows asymptotically, the size of $\mathcal{G}_\mathcal{C}$ also grows asymptotically.*

Proof. The probability of a gene tree to exhibit coverage pattern $C$ is $p(C)$, which by definition of $\mathcal{C}_\mathcal{M}$, is strictly positive. Thus, when the number of gene trees grows asymptotically, the number of gene trees with coverage pattern $C$ also grows asymptotically.

Lemma 1.2. *When the size of $\mathcal{G}_\mathcal{C}$ grows asymptotically, $S_{|C}$ is the species tree with leaf set $C$ that minimizes the global length $L(S_{|C}, \mathcal{G}_\mathcal{C})$.*

Proof. All gene trees from $\mathcal{G}_\mathcal{C}$ cover all species of $S_{|C}$ and the global length score is consistent under the multi-species coalescent (MSC) model for complete gene trees (Vachaspati and Warnow, 2015).

Lemma 1.3. *The global induced length of $S$ is equal to:*

$$L_p(S, \mathcal{G}) = \sum_{C \in \mathcal{C}_S} L(S_{|C}, \mathcal{G}_\mathcal{C}) \quad (1)$$

Proof.

$$L_p(S, \mathcal{G}) = \sum_{k=1}^{K} L(S_k, \{G_k\}) \quad (2)$$

$$= \sum_{k=1}^{K} \sum_{i \in L(G_k)} \sum_{j \in L(G_k)} 2^{-M_k(i,j)} D_k(i,j) \quad (3)$$

$$= \sum_{C \in \mathcal{C}_S} \sum_{k \in \mathcal{G}_\mathcal{C}} \sum_{i \in C} \sum_{j \in C} 2^{-M_k(i,j)} D_k(i,j) \quad (4)$$

$$= \sum_{C \in \mathcal{C}_S} L(S_{|C}, \mathcal{G}_\mathcal{C}) \quad (5)$$

Theorem 1.4. *When the size of $\mathcal{G}$ grows asymptotically, the global induced length of $\mathcal{G}$ is minimized by $S$.*

Proof. When the size of $\mathcal{G}$ grows asymptotically, each term of the sum in Eq. 1 is minimized by $S$ (Lemma 1.1 and Lemma 1.2). Thus, the global induced length is also minimized by $S$ (Lemma 1.3).

Theorem 1.4 proves that the the global induced length score is consistent under the multi-species coalescent model and under any random model of taxon deletion that is independent across partitions, provided that the gene trees have been correctly estimated.

© The Author 2022.



picture(0,0)(-35,0)(1,0)30 (0,-35)(0,1)30 picture picture(0,0)(35,0)(-1,0)30 (0,-35)(0,1)30 picture

"output" — 2022/7/22 — page 1 — #1

picture(0,0)(-35,0)(1,0)30 (0,35)(0,-1)30 picture picture(0,0)(35,0)(-1,0)30 (0,35)(0,-1)30 picture



#### 2 Tree search heuristic

In order to find a species tree with a 'good' score, we start from a given species tree topology and apply a tree search heuristic to further improve the score. The starting species tree can be either a random tree or a tree estimated with our own re-implementation of the ASTRID algorithm. We first briefly outline the FastME search algorithm (Lefort *et al.*, 2015), which is used in ASTRID, and then explain how we adapt it to optimize the global induced length.

FastME takes as input a distance matrix $D$ and applies subtree prune and regraft (SPR) moves to further improve it with respect to the minimum global length. At every step, FastME first precomputes the so-called *average distance* between every pair of subtrees in the current species tree in $O(N^2)$ time where $N$ is the number of species. After this precomputation step, computing the global length of any new species tree that can be obtained via an SPR move can be conducted in constant time. FastME thus identifies the optimal SPR move in $O(N^2)$ time (there are $O(N^2)$ possible SPR moves) and applies it. We call this optimial SPR move identification step *FastMEEval*. This procedure is repeated until no species tree with a lower minimum global length can be found. Assuming that $D$ is given, the entire search runs in $O(pN^2)$ time, where $p$ is the number of iterations required until convergence. The algorithm is described in more detail in the original paper (Lefort *et al.*, 2015) and is also not guaranteed to find the globally optimal tree.

Given the current species tree topology $S$, Asteroid computes for each gene tree $G_k$ the species tree $S_k$ induced by $S$ and $L(G_k)$. Then, for every gene $k$, it applies the FastMEEval procedure to $S_k$ and to its corresponding gene internode distance matrix $D_k$, in order to evaluate every SPR move that can be applied to $S_k$. This can be done in $O(\sum_{G_k \in \mathcal{G}} |G_k|^2)$ time. Finally, Asteroid can evaluate every SPR move that can be applied to $S$ by summing over the costs of all corresponding SPR moves in the induced species trees. Evaluating all moves requires $O(KN^2)$ time. For substantial proportions of missing genes ($> 50\%$), $O(KN^2) \gg \sum_{G_k \in \mathcal{G}} |G_k|^2$. Hence, this step is the most computationally expensive in practice. For this reason, we limit the radius of the SPR moves we evaluate: for a given pruning node, we explore the candidate re-grafting nodes by recursively traversing the remaining part of the tree, but stop the recursion after $q$ recursive calls without improvement. In our implementation, we set $q := 3$, and add an additional step with $q = \infty$ if no better tree can be found. We explored appropriate values for $q$ empirically on various empirical datasets to obtain the best trade-off between runtime and accuracy.

In ASTRID, most of the runtime is spent in evaluating the input distance matrix $D$. In Asteroid however, the largest proportion of runtime is spent in evaluating the SPR moves. As a result, the overall runtime is approximately linear to the number of iterations $p$ required for the algorithm to converge. For this reason, instead of only applying the optimal SPR move, Asteroid applies several moves simultaneously. Our algorithm maintains a list of all moves with a positive predicted score and sorts them in ascending order. Then, it sequentially applies all these moves, but omits the moves that conflict with previously applied moves. Two SPR moves are conflicting if they have the same pruning or regrafting nodes. The resulting global length is only computed after *all* of these moves have been applied. Note that, the prediction of an SPR move score is exact only if this SPR move is exclusively applied to the tree. Thus, there is not guarantee that applying several SPR moves with a positive *predicted* score will result in a species tree with a better score. If the new species tree has a worse score than the preceding tree, we rollback to the preceding tree and only apply the best SPR move instead.

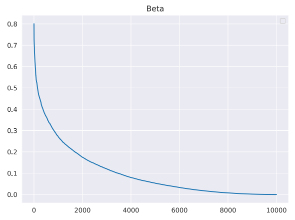


**Fig. 1.** Very high proportion of missing data: $B^*(0.1, 5)$

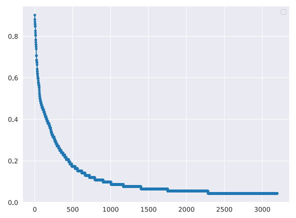

**Fig. 2.** Life92 per-gene coverage

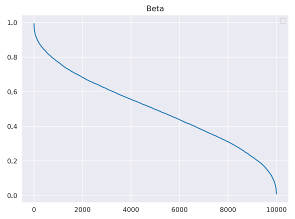


**Fig. 3.** Medium proportion of missing data: $B^*(0.5, 5)$

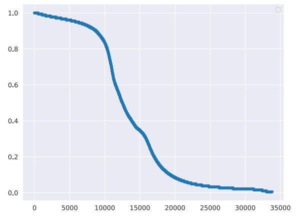

**Fig. 4.** Vertebrates179 per-gene coverage

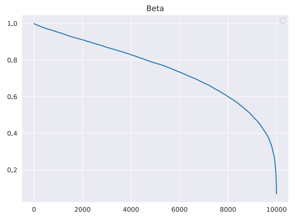


**Fig. 5.** Low proportion of missing data: $B^*(0.75, 5)$

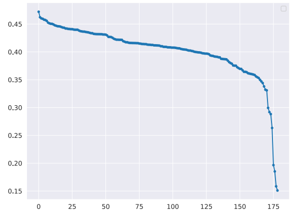

**Fig. 6.** Vertebrates179 per-species coverage

**Fig. 7.** Density of the reparametrized Beta distributions for $\mu \in \{0.1, 0.5, 0.75\}$ and $\theta = 5$, and empirical distributions of gene occupancy (per-family and per-species). The datasets Life92 and Vertebrates179 are described in the Empirical datasets subsection of the main paper.

#### 3 Missing data distribution shapes

##### 3.1 SimPhy parameters

We detail the parameters used to generated the simulated datasets with SimPhy in Table. 1 and Table. 2

#### 4 Building the reference vertebrate tree

In order to build the reference vertebrate tree, we started from the NCBI taxonomy tree, and then collapsed the branches between the following clades:

- Serinus Canaria, Geospiza Forti, Taenioygia Guttata (birds)
- Rhinolophus Ferrumequinum, Myotis Lucifugus, Pteropus Vampyrus (bats)
- Bos Mutus, Bos Taurus, Bison Bison
- Saimiri Boliviensis, Callithrix Jacchus, Aotus Nancymaae (new world monkeys)
- Ambassidae, Pomacentridae, and Cichliformes (fishes)
- Dicentrarchus Labrax and Larimichthys crocea (fishes)

picture(0,0)(-35,0)(1,0)30 (0,-35)(0,1)30 picture picture(0,0)(35,0)(-1,0)30 (0,-35)(0,1)30 picture

"output" — 2022/7/22 — page 2 — #2

| Default parameters | |
|---|---|
| Speciation and extinction rates | $5 \times 10^{-9}$ and $4.9 \times 10^{-9}$ |
| Number of gene families | 100 |
| Number of species | 50 |
| Population size | 10 |
| Per-species deletion probability | 0 |
| Per-family deletion probability | 0 |
| Species tree height | $Log\text{-}\mathcal{N}(21.25, 0.2)$ |
| Global substitution rate | $Log\text{-}\mathcal{N}(-21.9, 0.1)$ |
| Lineage specific rate gamma shape | $Log\text{-}\mathcal{N}(1.5, 1)$ |
| Family specific rate gamma shape | $Log\text{-}\mathcal{N}(1.551533, 0.6931472)$ |
| Gene tree branch specific rate gamma shape | $Log\text{-}\mathcal{N}(1.5, 1)$ |
| Sequence length | $\nu \times Log\text{-}\mathcal{N}(0, 0.25), \nu = 100(e^{-\frac{0.25^2}{2}})$ |
| Sequence base frequencies | Dirichlet(A=36,C=26,G=28,T=32) |
| Sequence transition rates | Dirichlet(TC=16,TA=3,TG=5, CA=5,CG=6,AG=15) |
| Seed | $[3000, 3050[$ |
| **Varying parameters** | |
| Per-species deletion probability | $B^*(\mu_s, 0.5), \mu_s \in \{0.4, 0.5, 0.6, 0.7, 0.8\}$ |
| Per-gene deletion probability | $B^*(\mu_f, 0.5), \mu_f \in \{0.4, 0.5, 0.6, 0.7, 0.8\}$ |
| Per-species and per-gene deletion probabilities | $(B^*(\mu, 0.5), B^*(\mu, 0.5)), \mu \in \{0.2, 0.3, 0.4, 0.5, 0.6\}$ |

Table 1. SimPhy parameters to study the effect of the missing data distribution (without ILS).

| Default parameters | |
|---|---|
| Speciation and extinction rates | $5 \times 10^{-9}$ and $4.9 \times 10^{-9}$ |
| Number of gene families | 1000 |
| Number of species | 50 |
| Population size | $5 * 10^7$ |
| Per-species deletion probability | $B^*(0.6, 0.5)$ |
| Per-family deletion probability | $B^*(0.6, 0.5)$ |
| Species tree height | $Log\text{-}\mathcal{N}(21.25, 0.2)$ |
| Global substitution rate | $Log\text{-}\mathcal{N}(-21.9, 0.1)$ |
| Lineage specific rate gamma shape | $Log\text{-}\mathcal{N}(1.5, 1)$ |
| Family specific rate gamma shape | $Log\text{-}\mathcal{N}(1.551533, 0.6931472)$ |
| Gene tree branch specific rate gamma shape | $Log\text{-}\mathcal{N}(1.5, 1)$ |
| Sequence length | $\nu \times Log\text{-}\mathcal{N}(0, 0.25), \nu = 100(e^{-\frac{0.25^2}{2}})$ |
| Sequence base frequencies | Dirichlet(A=36,C=26,G=28,T=32) |
| Sequence transition rates | Dirichlet(TC=16,TA=3,TG=5,CA=5,CG=6,AG=15) |
| Seed | $[3000, 3050[$ |
| **Varying parameters** | |
| Per-species and per-gene deletion probabilities | $(B^*(\mu, 0.5), B^*(\mu, 0.5))$,<br>$\mu \in \{0.5, 0.55, 0.6, 0.65, 0.7, 0.75\}$ |
| population size | 10, $5 * 10^7$, $10^8$, $5 * 10^8$, $10^9$ |
| Number of gene families | 250, 500, 1000, 2000 |
| Gene tree branch length scaler | 0.05, 0.1, 1.0, 10.0, 100.0, 200.0 |
| Average sequence length | 50, 100, 200, 500 |
| Number of species | 25, 50, 75, 100, 125, 150 |

Table 2. SimPhy parameters to study the effect of the remaining parameters.

#### 5 Identifying terraces

Datasets with high levels of missing data are more likely to produce *terraces*, that is, sets of distinct species trees that have exactly the same analytical score (Sanderson *et al.*, 2015). This is because, in presence of missing data, several species trees might belong to the same *stand*, defined as the set of species trees that induce the same per-gene trees (the set of trees $S_k = S_{|L(G_k)}$ as defined in our Methods section). Trees that belong to the same stand also belong to the same terrace with respect to the optimization criteria used in Asteroid, FastRFS, and ASTRAL-III. Note that, this is not true for the global length criteria used in ASTRID. We say that a dataset whose best-scoring tree belong to a terrace with strictly more than one tree is *indecisive*. Identifying indecisive datasets is crucial, because most

picture(0,0)(-35,0)(1,0)30 (0,35)(0,-1)30 picture
picture(0,0)(35,0)(-1,0)30 (0,35)(0,-1)30 picture



tools only return one single tree, even when there exist many trees that are equally good.

In order to identify indecisive datasets, we ran Gentrius, a tool implemented in IQTREE-2 Minh *et al.* (2020), release 2.2.0. Gentrius takes as input a presence-absence matrix whose rows are the genes and whose columns are the species. The elements are set to 1 if the gene covers the species, and 0 otherwise. Then, for a given input species tree, Gentrius returns the size of its stand. We ran Gentrius on all empirical datasets, using the Asteroid tree as input.

According to Gentrius, none of the trees inferred from the Life92-single and Life92-disco datasets belong to a terrace with more than one tree, and both datasets are thus decisive. However, the two filtered Vertebrates179 datasets with the highest proportion of missing data are indecisive: With $\tau = 0.8$, the Asteroid tree belong to a terrace of three trees, and with $\tau = 0.9$, to a terrace of more than 1,000,000 trees.

picture(0,0)(-35,0)(1,0)30 (0,-35)(0,1)30 picture
picture(0,0)(35,0)(-1,0)30 (0,-35)(0,1)30 picture
"output" — 2022/7/22 — page 4 — #4
